# Tomato brown rugose fruit virus and pepino mosaic virus differentially modulate disease severity of bacterial pith necrosis and bacterial canker in tomato

**DOI:** 10.64898/2026.07.29.741454

**Authors:** Evanson Ngugi, Ilana Bekelman, Nir Berholz, Lior Avraham, Sigal Perets, Edward Belausov, Maya Bar, Aviv Dombrovsky, Doron Teper

## Abstract

Tomato worldwide production is increasingly challenged by complex disease outbreaks involving multiple interacting pathogens. In Israel, recent years have seen a marked rise in vascular collapse symptoms in greenhouse-grown tomatoes, coinciding with the widespread emergence of tomato brown rugose fruit virus (ToBRFV) and pepino mosaic virus (PepMV). Here, we investigated the bacterial and viral agents associated with these outbreaks and examined how viral infection influences the development and severity of bacterial diseases that cause vascular collapse. Surveys conducted between 2021 and 2026 revealed that tomato pith necrosis outbreaks were associated with a diverse bacterial community dominated by members of the *Pseudomonadales* and *Enterobacterales*, while bacterial canker outbreaks were exclusively linked to *Clavibacter michiganensis*. Multilocus sequence analysis showed that pith necrosis–associated *Pseudomonas* isolates clustered primarily within the *P. syringae*, and *P. corrugata* phylogroups. Pathogenicity assays demonstrated that only a subset of pith necrosis-associated bacteria, *P. mediterranea*, *P. capsici*, *P. viridiflava*, and *Xanthomonas euvesicatoria* pv*. perforans*, induced pith necrosis under controlled conditions, with high variability in symptom severity. Co-inoculation experiments showed that ToBRFV- and PepMV-infected plants exhibited a 40-100% increase in lesion size following inoculation with pith necrosis-associated bacteria, without a corresponding increase in bacterial colonization, whereas the same viral infections attenuated wilt symptoms caused by *C. michiganensis*. Together, our findings demonstrate that endemic viral infections differentially modulate bacterial disease outcomes, either exacerbating or attenuating symptoms depending on the pathogen. These results highlight the importance of multi-pathogen interactions in disease severity and have important implications for tomato disease management.

## Introduction

Worldwide tomato (*Solanum lycopersicum*) production is significantly affected by microbial pests, including viruses, fungi, and bacteria, causing substantial yield losses and economic damage to the global tomato industry [1]. Over the past decade, the tomato sector has suffered major losses due to the emergence of tomato brown rugose fruit virus (ToBRFV, *Tobamovirus fructirugosum*) [2]. ToBRFV, a highly infectious tobamovirus, was first reported in tomato greenhouses in Jordan in the spring of 2015 [3], with retrospective analyses tracing its earliest outbreaks to Israel in 2014, where it overcame the key resistance genes *Tm-2* and *Tm-2^2^* [4]. Since then, ToBRFV has spread to nearly all tomato-producing countries worldwide, resulting in severe and widespread yield losses [5]. ToBRFV is primarily transmitted in nethouses and greenhouses through mechanical contact during routine agricultural practices such as pruning, trellising, and spraying, and over long distances via contaminated seeds and fruits [5]. Disease symptoms include brown, rough lesions on leaves, sepals, and stems, as well as yellow discoloration on ripe fruits [4,6]. Infected leaves often exhibit mosaic patterns accompanied by narrowing and mottling [2]. In Israel, ToBRFV is frequently associated with co-infection by a second viral pathogen, pepino mosaic virus (PepMV, *Potexvirus pepini*) [7]. PepMV is a potexvirus that infects several solanaceous crops and causes fruit marbling, severe mosaic symptoms, leaf bubbling and distortion, and, as recently reported, necrosis and growth stunting [8]. Recent surveys indicate that both viruses are present in most commercial tomato greenhouses in the country [7] and are also maintained in wild Mediterranean plant populations [9,10]. Notably, a synergistic interaction between the two viruses has been demonstrated, in which ToBRFV infection increases host susceptibility to PepMV and enhances PepMV replication, thereby exacerbating PepMV-associated yield losses [7].

In addition to viral pathogens, both global and Israeli tomato production are affected by bacterial diseases, particularly bacterial canker and pith necrosis, which are prevalent throughout the country. Tomato bacterial canker is caused by the actinobacterium *Clavibacter michiganensis*, which systemically colonizes host xylem vessels, leading to stem cankers, wilting, leaf blotching, and vascular collapse [11]. *C*. *michiganensis* has been reported in most tomato-growing regions worldwide and has caused numerous outbreaks that threaten tomato production [12]. As a result, *C*. *michiganensis* has been, or is currently, under eradication programs in the European Union and is considered a quarantine pest in many countries [12]. Israel experienced several severe outbreaks of bacterial canker in the early 2000s that nearly devastated the tomato industry. In response, a large-scale integrated pest management (IPM)– based control program was implemented, which largely contained the pathogen [13]. However, although its impact has been substantially reduced, *C*. *michiganensis* remains endemic throughout the country [14]. Tomato pith necrosis is a stem disorder characterized by internal browning, collapse, and hollowing of the stem pith, leading to wilting, reduced plant vigor, adventitious root formation along the stem, and, in severe cases, plant collapse [15]. The disease predominantly affects greenhouse-grown tomatoes under conditions of high humidity and vigorous vegetative growth, typically during fruit ripening, and often appears as patchy outbreaks within a greenhouse [15]. In contrast to bacterial canker, which is caused by a highly specialized pathogen and exhibits well-defined and distinctive symptoms, the etiology of pith necrosis is less clearly defined and has been associated with multiple bacterial taxa; accordingly, it is often described as a conditional or opportunistic disease. Initially, tomato pith necrosis was linked to *Pseudomonas corrugata*, which was later separated into *P. corrugata* and *P. mediterranea* [16]. Subsequently, additional bacterial species were reported in association with the disease, including several *Pseudomonas* species from the *P. fluorescens*, *P. syringae*, and *P. putida* phylogroups, as well as *Xanthomonas euvesicatoria* pv. *perforans*, *Enterobacter* spp., and soft-rot Pectobacteriaceae (SRP), such as *Pectobacterium* and *Dickeya* [17–19]. Due to its opportunistic nature and the large number of associated bacterial taxa, tomato pith necrosis remains poorly studied, and little mechanistic insight into disease development is currently available. In Israel, pith necrosis has been reported since the 1980s [20]; however, because of its relatively minor economic impact at the time, it was not subjected to in-depth investigation, and the associated bacterial populations remain poorly characterized. However, this has shifted in recent years, with bacterial vascular collapse emerging more frequently in greenhouse tomato production over the past decade. This disease, which displays pith necrosis–like symptoms, has resulted in significant yield losses. Notably, these phenomena have frequently been reported in greenhouses affected by viral outbreaks, suggesting that interactions among multiple pathogenic agents may contribute to disease development. Notably, the interaction between ToBRFV and PepMV and bacterial vascular diseases has yet to be studied, and it remains unknown whether these endemic viral infections alter the etiology and severity of bacterial vascular diseases in tomato production systems.

During this study, we characterized bacterial and viral agents associated with tomato vascular collapse, identify which bacteria are causal agents of with tomato vascular collapse under controlled conditions, and evaluated the influence of viral infection on the development and severity of bacterial diseases in tomato.

## Materials and Methods

### Plant material and growth conditions

Tomato plants (cv. Ikram) were obtained as seedlings at the two-leaf stage from from Hishtil nursery (https://www.hishtil.com/). Seedlings were transplanted into GREEN 90 potting mix containing slow-release fertilizer (Pelemix GREEN; https://www.evenari.co.il/html5/prolookup.taf?&_id=23981&did=9414&title=%20%FA%F2%F8%E5%E1%FA%20%F9%FA%E9%EC%E4%20green%2090) and transferred to a temperature-controlled glasshouse maintained at 25 ± 2 °C. Plants were grown under natural light conditions, with relative humidity ranging between 60% and 90% (average 75%). Irrigation was performed manually on a daily basis.

### Sample collection, viral diagnostics and bacterial isolation

Tomato plant samples showing symptoms of vascular collapse were collected from commercial tomato greenhouses in Israel between 2021 and 2026. Samples were visually assessed for symptoms consistent with pith necrosis or bacterial canker and were sent to an external diagnostic laboratory for testing for common fungal and oomycete vascular pathogens, including *Fusarium* spp., *Pythium* spp., and *Sclerotium rolfsii*. When no clear fungal causal agent was identified, samples were subsequently tested for the presence of ToBRFV and PepMV and used for the isolation of putative bacterial pathogens.

### Indirect Enzyme-Linked Immunosorbent Assay (ELISA) based viral detection and isolation of stem-associated bacteria

For viral identification plant samples were subjected to an indirect ELISA test [21]. Each sample had a technical duplicate repeat, as well as a buffer control and positive and negative controls. For ToBRFV and PepMV detection, two leaflets were collected from each plant exhibiting pith necrosis or bacterial canker symptoms and subjected to virus detection by ELISA using ToBRFV and PepMV-specific antibodies, as described previously [7]. The developed color was recorded using ELISA reader (Thermo Fisher Multiskan FC) at 405 nm and 620 nm.

For bacterial isolation, plant samples were washed thoroughly and surface-sterilized using 75% ethanol. Bacterial isolation was performed by excising internal stem tissues from symptomatic plants, avoiding heavily necrotic regions and preferentially selecting sections showing mild symptoms indicative of early disease stages. Excised tissues were placed in 1.5-mL microcentrifuge tubes containing 250 µL of double-distilled water and macerated using a grinder fitted with a 1.5-mL pellet pestle to generate a suspension. The suspension was then brought to a final volume of 1 mL with double-distilled water and used to prepare five ten-fold serial dilutions. Aliquots (10 µL) from each dilution were spotted onto Luria broth (LB) agar plates and incubated at 28 °C. Bacterial colonies were monitored for 1–3 days. In cases of high bacterial load, where colonies were detectable at dilutions of at least 10 ³ and a single bacterial morphotype predominated, individual colonies were streaked onto fresh LB agar plates and incubated under the same conditions to obtain pure cultures. Selected colonies were preserved as glycerol stocks by suspending cells in 25% glycerol in 1.5-mL microcentrifuge tubes and storing them at −80 °C. Corresponding plates were stored at 4 °C for short-term use in subsequent experiments.

### Bacterial identification

After isolation, dominant bacterial colonies from each sample were differentiated based on colony morphology and purified by quadrant streaking to obtain single-colony isolates. For samples exhibiting pith necrosis–like symptoms, which were associated with diverse bacterial taxa, representative colonies were initially identified by PCR amplification and sequencing of the 16S rRNA gene using universal bacterial 16S rRNA gene primers (Table S1). Bacterial species were assigned based on sequence similarity using NCBI nBLAST searches (https://blast.ncbi.nlm.nih.gov/Blast.cgi). When 16S rRNA gene sequencing did not reliably resolve closely related taxa, the *gyrB* gene was PCR-amplified and sequenced using primer sets designed to target members of the orders *Pseudomonadales*, *Xanthomonadales*, *Enterobacterales*, and *Micrococcales* (Table S1). For samples displaying bacterial canker symptoms, bacterial colonies were visually screened for morphology characteristic of *C*. *michiganensis* and subsequently confirmed by diagnostic PCR using primers targeting the *chpC* gene within the *C*. *michiganensis chp/tomA* pathogenicity island [22] and/or by amplification and sequencing of the *gyrB* gene using *Micrococcales*-specific primers.

### Multilocus Sequence Analysis

The phylogenetic lineage of pith necrosis–associated *Pseudomonas* species was assessed using multilocus sequence analysis (MLSA). Primer sets designed to amplify three housekeeping genes (*gyrB*, *rpoB*, and *recA*) from members of the order *Pseudomonadales* (Table S1) were used for PCR amplification and sequencing of all *Pseudomonas* isolates included in the analysis. For each isolate, the three gene sequences were concatenated in a fixed gene order to generate a combined multilocus sequence. Reference sequences corresponding to the concatenated loci were retrieved from publicly available complete genome sequences of *Pseudomonas* species, with *Ralstonia solanacearum* included as an outgroup, using the NCBI database. Multilocus sequence alignments and phylogenetic analyses were performed using MEGA 11 software (https://www.megasoftware.net/). Phylogenetic trees were constructed using the maximum likelihood method with 200 bootstrap replicates to assess node support.

### ToBRFV and PepMV Viral inoculation and detection by ELISA

Leaf extracts of *Nicotiana tabacum*, *Datura stramonium* and tomato plants, previously inoculated with ToBRFV, PepMV, and ToBRFV + PepMV, respectively were used as viral inoculum for the experiments. Prior to infection, viral inoculum was confirmed with ELISA using ToBRFV and PepMV specific antibodies as as described by [7]. Sap mechanical inoculation, were done by gently rubbing leaflets from the third and fourth leaves of four-leaf stage tomato plants cv. Ikram with viral inoculum supplemented with carborundum to enable symplastic delivery as described previously [23]. Non-infected plant extract was used as a “mock” negative control for each viral inoculation experiment. After viral inoculations, plants were then kept in a temperature-controlled glasshouse at 25 ± 2 °C under natural light. Viral presence was estimated three to four weeks post inoculation by taking two leaflets from the youngest leaf in each plant and taking at least three representative plants per treatment. The presence of both viruses was tested for each sample type (no virus control, ToBRFV alone, PepMV alone, ToBRFV + PepMV) to confirm viral infection and to verify that undesired viral cross contamination did not occur.

### Plant inoculation with pith necrosis–associated bacteria and assessment of lesion size and bacterial populations

For plant inoculation, pith necrosis-associated bacteria were streaked from glycerol stocks on fresh LB agar plates and incubated for 12-72 h (depending on growth rate of each bacterium) at 28° C. Freshly grown bacterial cultures were pooled from the plate using a sterile toothpick and directly stabbed into the stem section between the cotyledons of four-leaf-stage tomato plants cv. Ikram. For negative control plant were “mock” inoculated with sterile toothpick alone. In co-inoculation experiments, bacteria were inoculated into plants 24 h after the initial viral or mock inoculations. After bacterial inoculations, plants were then kept in a temperature-controlled glasshouse at 25 ± 2 °C under natural light. The glasshouse was divided into four sections using plastic barriers to prevent viral transmission between plants. Each section was used separately for a specific viral inoculation treatment in each experiment, while mock- and bacteria-inoculated plants within each viral treatment were randomized. Each experiment included 6–17 biological replicates per treatment and was repeated five times for ToBRFV + PepMV co-inoculations or twice for independent ToBRFV or PepMV inoculations. To minimize positional bias within the greenhouse, the assigned sections for viral inoculations were rotated between experimental repeats. Bacterial virulence was assessed one month after inoculation by quantifying disease symptoms and stem bacterial populations. For assessment of symptoms, infected stems were horizontally split into two for visualization of inner brown lesions of the pith tissues. Split stems were than photographed and lesion surface areas quantified using the ImageJ image analysis software (https://imagej.net/ij/) by measuring the brown-pigmented lesion area in each plant. For bacterial populations quantification, 1-3 mm horizontal stem tissues were excised from the site of infection and 2 cm above the point of infection. These tissues were weighed, surface sterilized and macerated as described in ’sample collection, viral diagnostics and bacterial isolation’ and subjected to 10-fold serial dilutions, spotted on LB agar, plated and incubated at 28° C for 24-72 h until single colonies appear (depends on the bacteria). Colony forming units (CFU) were counted in the lowest dilution of which single colonies could be separated from each other and bacteria titre per each sample was calculated as CFU/gram stem.

### Plant inoculation with *Clavibacter michiganensis* and assessment of wilt symptoms and bacterial populations

Inoculation and disease assessment of bacterial canker were conducted as described previously [24]. *C*. *michiganensis* strain NCPPB382 was streaked from glycerol stocks onto fresh LB agar plates and incubated for 72 h. Bacteria were collected from freshly grown plates into a 1.5 ml centrifuge tube containing sterile double-distilled water (DDW) and diluted to approximately 5 × 10 CFU/ml (OD600 = 0.1) in 1 ml DDW. Sterile toothpicks were incubated in the diluted bacterial suspension for 5–10 min and then used to puncture the stem between the cotyledons of four-leaf-stage tomato plants cv. Ikram. For negative controls, plants were mock-inoculated using sterile toothpicks only. In co-inoculation experiments, bacterial inoculation was performed 24 h after the initial viral or mock inoculations. Plants were maintained in a temperature-controlled glasshouse at 25 ± 2 °C under natural light. The glasshouse was divided into four sections using plastic barriers to prevent viral transmission between plants. Each section was used separately for a specific viral inoculation treatment in each experiment, while mock- and *C*. *michiganensis*-inoculated plants within each viral treatment were randomized. Each experiment included 4–10 biological replicates per treatment and was repeated five times for ToBRFV + PepMV co-inoculations or twice for independent ToBRFV or PepMV inoculations. To minimize positional bias within the greenhouse, the assigned sections for viral inoculations were rotated between experimental repeats. Bacterial virulence was assessed three weeks post-inoculation by evaluating disease symptoms and quantifying stem bacterial populations. Wilt symptoms were visually monitored in each plant and assigned a virulence score based on the percentage of wilted leaves as follows: 0 = no wilt; 1 = 1–20%; 2 = 21–60%; 3 = 61–100%. Stem bacterial populations were quantified at the site of inoculation and 5 cm above the inoculation site, as described above.

### Bacterial transformation and visualization of GFP-labelled bacteria in planta

The broad-host-range plasmid pKT-Kn [25], a pVSP61 derivative constitutively expressing the green fluorescent protein (GFP) gene under the control of the *trp* promoter, was introduced into *P. mediterranea* (Pmed, G51), *P. viridiflava* (Pvir, G84), and *P. capsici* (Pcap, G143) by electroporation as described previously [26]. Transformed clones were selected on LB agar supplemented with 50 µg/ml kanamycin, and GFP expression was confirmed by fluorescence using a stereomicroscope equipped with a GFP filter. To monitor tissue localization and bacterial movement *in planta*, tomato plants were inoculated with each GFP-labelled bacterial clone. Bacterial localization at the inoculation site and at 1.5 cm and 3 cm above the inoculation site was examined 14 days post-inoculation (dpi). Horizontal and vertical stem cross-sections were excised from infected plants and visualized using a Leica SP8 laser scanning confocal microscope with GFP excitation and emission settings at the Volcani Institute, microscopy unit. The presence of GFP-labelled bacteria was assessed in pith parenchyma tissues and vascular bundles.

### Data visualization, and statistical analyses

Prism (GraphPad) and Microsoft Excel were used for statistical analysis and data visualization throughout this study. All experiments were independently repeated two to five times, with the number of experiments and the number of biological replicates per experiment specified in the corresponding figure legends. Continuous data, such as pith necrosis lesion size and bacterial population measurements, were first assessed for normality using the Shapiro–Wilk test. As most datasets were non-normally distributed, differences between groups were analyzed using the non-parametric Mann–Whitney U test. In experiments involving multiple bacterial causal agents in parallel assays, p-values were adjusted for multiple comparisons using the Benjamini– Hochberg (BH) procedure. For bacterial canker disease severity scores, results were presented as stacked bar charts showing score distributions. Statistical differences between treatments were assessed using chi-square tests relative to the no-virus control.

## Results

### Bacterial and viral profiles of greenhouse tomato disease outbreaks in Israel

A significant increase in outbreaks of bacterial diseases in greenhouse tomatoes in Israel has been reported over the past five years. We aimed to determine whether these outbreaks coincide with infections by ToBRFV and PepMV, which have become prevalent in Israel during this period, and to identify the bacterial agents associated with these diseases. To this end, samples of diseased plants were collected from tomato greenhouses across Israel that experienced bacterial outbreaks between 2021 and 2026 (Fig. 1A). Samples were obtained by plant protection and vegetable crop extension agents following reports from growers and were used both for the isolation of putative bacterial causal agents and for the detection of ToBRFV and PepMV. Most outbreaks were reported in greenhouses located in the northwestern Negev and Sharon plain regions (Fig. 1). In addition, bacterial isolates from tomato pith necrosis outbreaks during 2014–2015, collected by Dr. Shulamit Manulis-Sasson (personal communication), were included in the analysis. The majority of bacterial outbreaks reported nationwide exhibited classic pith necrosis symptoms, characterized by plant collapse and internal stem rot with dark brown pigmentation throughout the main stem of fruiting plants (Fig. 1B). Several outbreaks of bacterial canker were also reported in the northwestern Negev region, characterized by wilting, leaf scorch, stem and petiole cankers, and vascular collapse (Fig. 1C). In addition to bacterial symptoms, most greenhouse tomato crops either exhibited viral disease symptoms or had been reported to be infected with ToBRFV and/or PepMV within the year preceding sample collection. This was confirmed by diagnostic testing, which showed that most samples were positive for at least one of the viruses, and that the majority were positive for both (Table 1). After isolation from infected tissues, single bacterial colonies that heavily colonized symptomatic tissues were subjected to molecular identification, using species-specific PCR for bacterial canker isolates or amplicon sequencing for pith necrosis isolates. As expected, all plants exhibiting bacterial canker were heavily infected with *C. michiganensis*, which was identified based on colony morphology and diagnostic PCR (Table 1). In contrast, bacteria isolated from plants showing pith necrosis symptoms were considerably more diverse and displayed clearly distinct colony morphologies among isolates. Nevertheless, the orders *Pseudomonadales* and *Enterobacterales* were overrepresented within this group and included multiple bacterial species previously implicated in pith necrosis, including *Pseudomonas mediterranea*, *P. viridiflava*, *P. capsici*, members of the *P. fluorescens* group, *Enterobacter* spp., and *Pectobacterium* spp. (Table 1). In addition, members of the genera *Xanthomonas* and *Curtobacterium*, which include several plant-pathogenic species, were also among the pith necrosis–associated isolates (Table 1). These data demonstrate that while bacterial canker incidents were exclusively associated with *C. michiganensis*, pith necrosis was associated with a diverse bacterial species in plants largely positive for ToBRFV and/or PepMV.

**Fig. 1.**
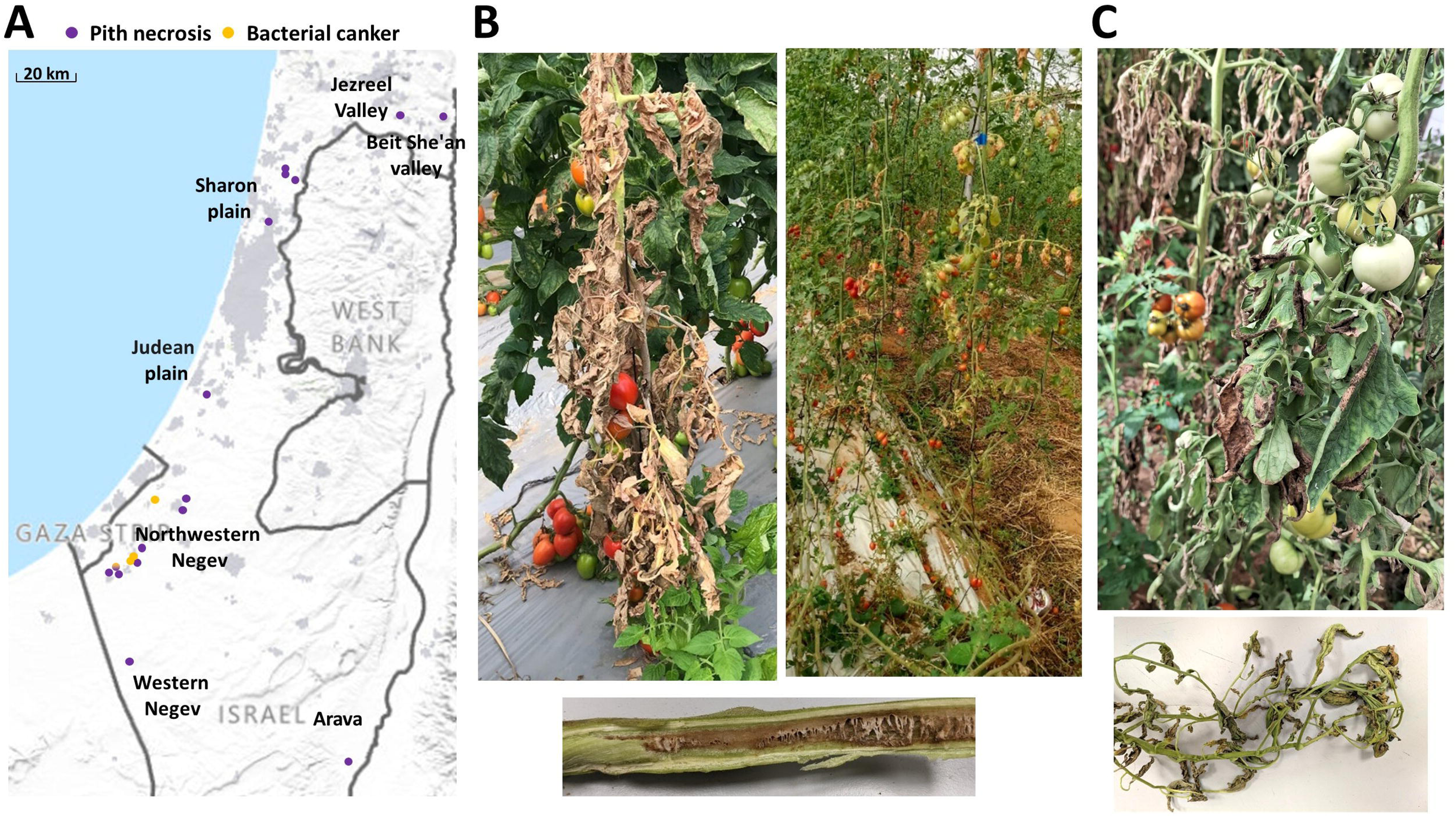
Geographic distribution and symptoms of tomato pith necrosis and bacterial canker outbreaks in Israel during 2021–2026. **(A)** Map showing the locations of pith necrosis (purple circles) and bacterial canker (orange circles) outbreaks in Israeli greenhouse tomato crops used for sample collection. Map source: U.S. Geological Survey (USGS), The National Map (https://www.usgs.gov/the-national-map). Data points and scale were added manually to the figure. Map indicates the geographic regions associated with disease outbreaks. **(B)** Upper panels show tomato plants with pith necrosis symptoms in commercial greenhouses, photographed in Prigan in the in the northwestern Negev (January 2022; left) and Beit Ezra in the Judean plain (April 2025; right). The lower panel shows stem pith necrosis in a Beit Ezra sample collected in 2025 and used for bacterial isolation. **(C)** Tomato plants displaying bacterial canker symptoms in Mivtachim in the northwestern Negev (November 2025). The lower panel shows leaf samples collected for bacterial isolation.

**Table 1.**
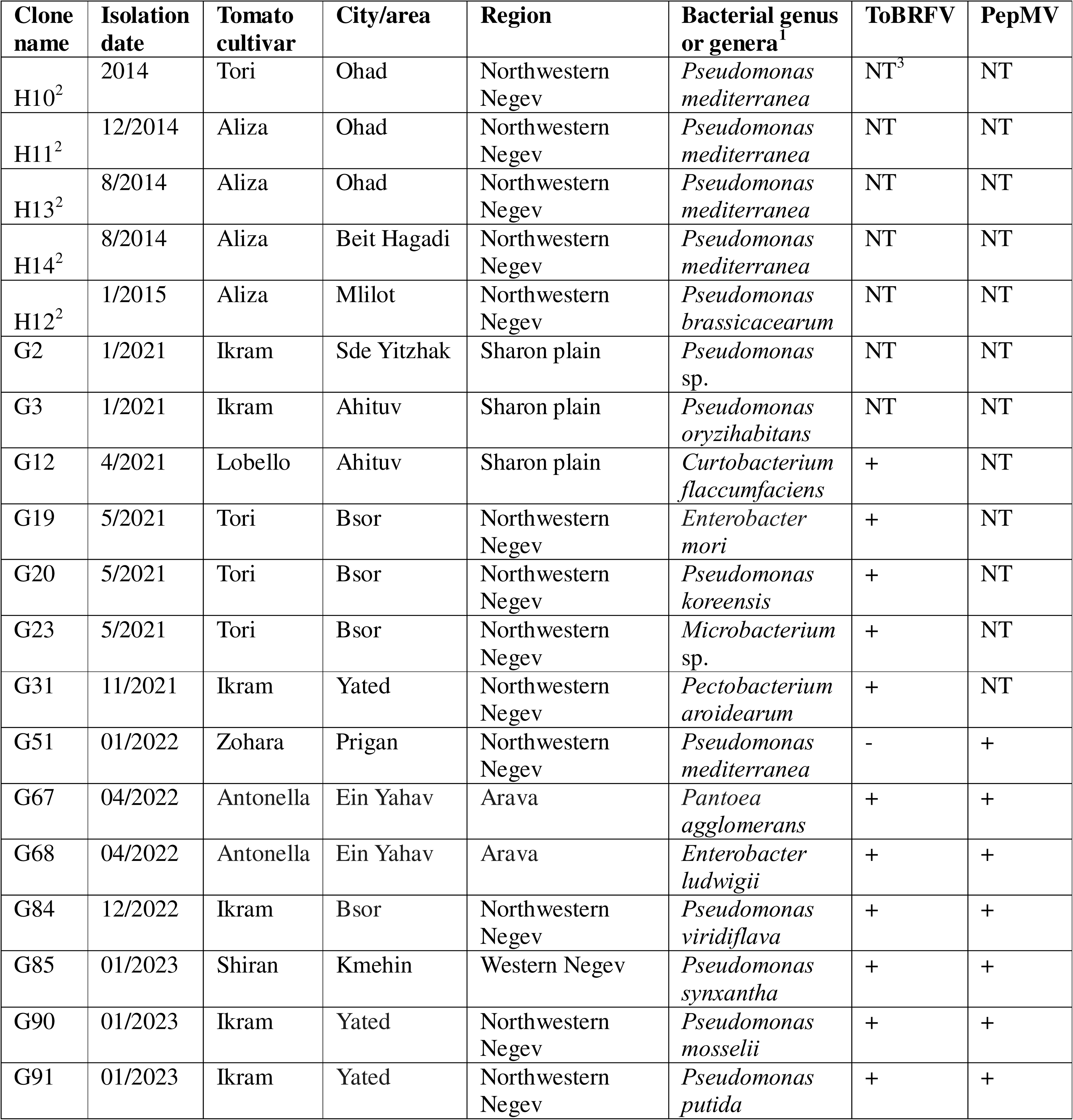
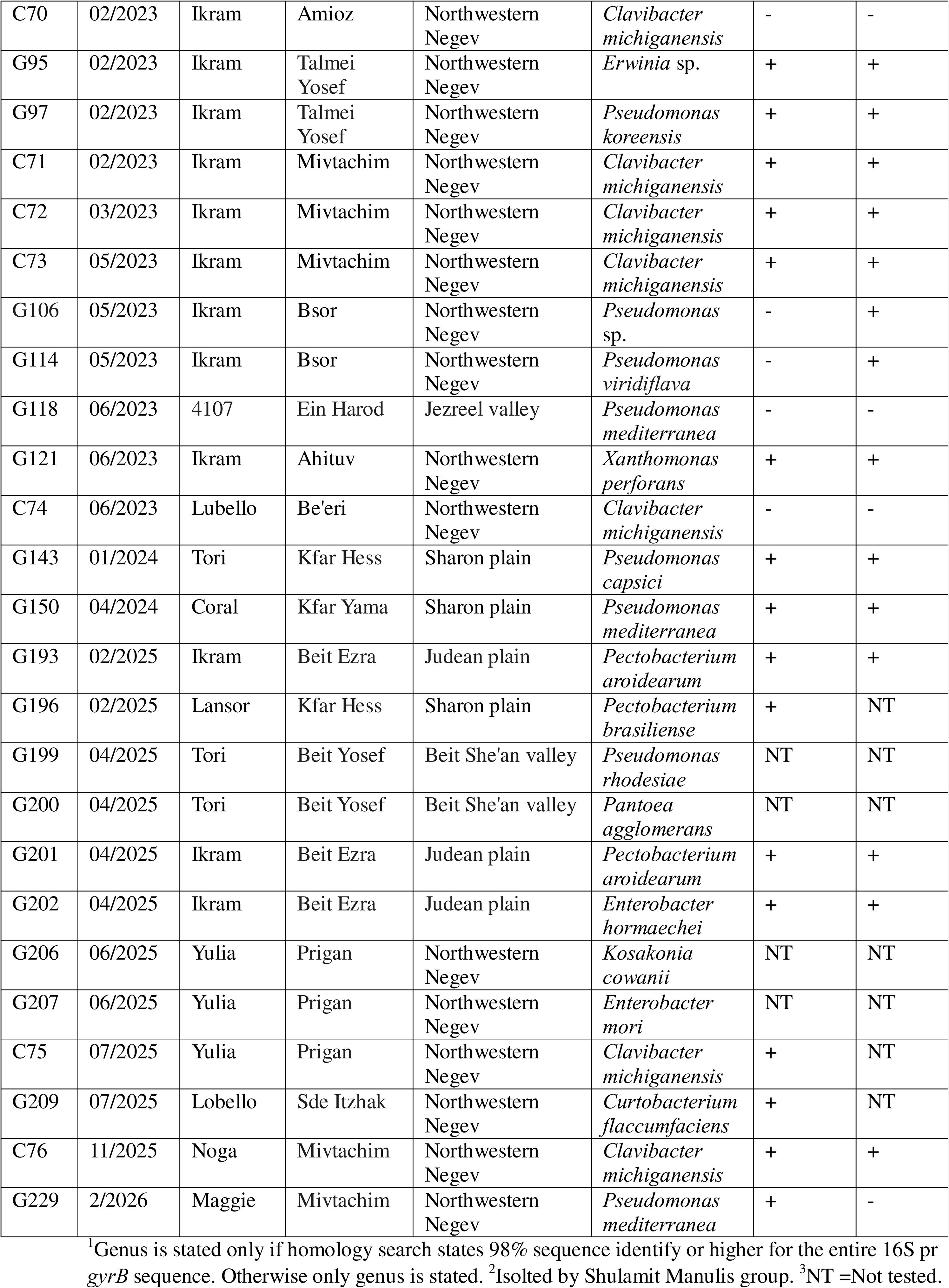
Presence of bacteria and viruses in tomato samples displaying vascular collapse in Israel.

### Phylogenetic lineages of pith necrosis-associated *Pseudomonadales*

*Pseudomonadales* constituted the dominant bacterial order among isolates from symptomatic tomato plants, consistent with their frequent association with pith necrosis. To achieve accurate species-level identification and assess the phyletic lineages of the isolates, a multilocus sequence analysis (MLSA) was performed and all isolates were resolved in a single maximum-likelihood phylogenetic tree together with appropriate reference genomes (Fig. 2). Most isolates clustered into two major phylogroups: the *P. corrugata* subgroup and the *P. syringae* group. Six isolates (H10, H11, H13, G51, G118, and G150) grouped tightly with *P. mediterranea* reference strains (DSM-16733 and B21-060), indicating limited intra-group diversity among these field isolates. Isolate G118 was nearly identical to *P. mediterranea* DSM-16733, while isolate H12, although positioned within the *P. corrugata* clade, branched separately and showed substantially lower sequence similarity to both *P. mediterranea* and *P. corrugata* reference genomes, suggesting a divergent lineage within this subgroup. Three isolates clustered within the *P. syringae* phylogroup: two isolates (G84 and G114) were identified as *P. viridiflava*, displaying high but non-clonal similarity to reference strains, while one isolate (G143) clustered with multiple *P. capsici* reference genomes, confirming its identity as *P. capsici*. Additional isolates were positioned within the *P. fluorescens* phylogroup: G85 and G106 clustered with *P. marginalis*, G97 and G20 grouped with *P. fluorescens* and *P. iransensis*, and G3 formed a distinct clade closely related to *P. oryzihabitans*. Collectively, the MLSA phylogeny reveals that tomato pith necrosis–associated *Pseudomonas* populations are dominated by closely related *P. mediterranea* strains, alongside genetically diverse members of the *P. syringae* and *P. fluorescens* phylogroups, including previously unreported species associations.

**Fig. 2.**
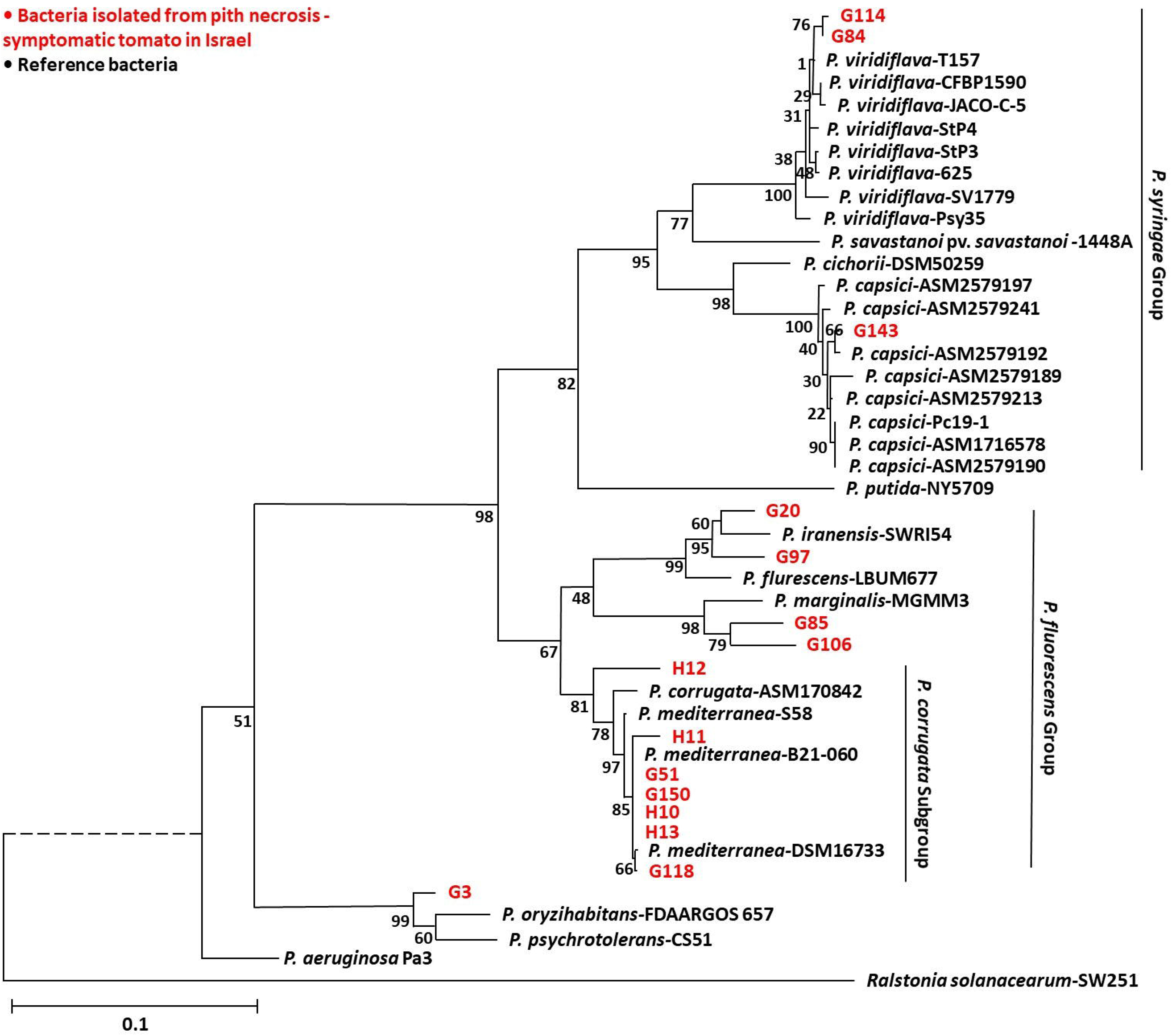
MLSA phylogeny of tomato pith necrosis isolates. Maximum-likelihood phylogenetic tree constructed from concatenated ***recA***, ***rpoB***, and ***gyrB*** gene sequences, depicting the taxonomic relationships among pith necrosis–associated bacterial isolates collected from multiple tomato greenhouses in Israel (highlighted in red) and reference strains obtained from the NCBI GenBank database. The phylogenetic analysis was performed using MEGA 11 with 200 bootstrap replicates. Bootstrap support values are shown at the branch nodes. Accession numbers of reference strain sequences are indicated next to the strain names and separated by a hyphen. The sequence of *Ralstonia solanacearum* strain UW251 was used as an outgroup.

### Virulence Assessment of Pith Necrosis-Associated Bacteria

After establishing the identity of pith necrosis–associated isolates, we evaluated their ability to cause disease and the severity of symptoms. Representative isolates from each identified species (one isolate when detected once and at least two when detected multiple times) were inoculated into ‘Ikram’ tomato plants using stab inoculation, and internal stem necrosis was assessed one-month post-inoculation. Most tested species failed to induce either localized or extensive necrosis under the tested conditions, suggesting that they are unlikely to be primary causal agents of pith necrosis in greenhouse-grown tomatoes, although their involvement in the disease under other conditions cannot be excluded. Although many field samples exhibiting classic pith necrosis symptoms contained *Pectobacterium* spp., inoculation with *P. aroidearum* and *P. brasiliense* resulted in severe soft rot that developed 48–96 h after inoculation (Fig. S1). This phenotype was more reminiscent of potato blackleg disease [27], leading to plant collapse rather than the inner browning typically associated with pith necrosis, possibly due to differences in growth conditions or infection mechanisms. In contrast, four species consistently induced pith necrosis with varying severity (Fig. S2). *P. viridiflava* (Pvir) caused weak to moderate lesions that rarely extended beyond the inoculation site, whereas *Xanthomonas euvesicatoria* pv. *perforans* (Xep) induced moderate lesions that expanded beyond the inoculation point but did not spread throughout the entire stem. *P. capsici* (Pcap) caused moderate to severe necrosis that frequently extended beyond the inoculation site and occasionally spread throughout the main stem, while *P. mediterranea* (Pmed) induced symptoms ranging from weak to severe, with lesions often extending beyond the inoculation site and, in some cases, throughout the entire stem. Notably, lesion size and disease severity showed high variability across biological and experimental replicates, ranging from minimal or no necrosis to extensive stem-wide lesions (Fig. S2). Next, we examined whether necrosis induced by pith necrosis–associated bacteria is liked to bacterial localization patterns by monitoring the tissue distribution of GFP-labeled Pcap and Pvir. Horizontal and vertical stem sections from inoculated plants were collected from the margins of necrotic areas, as well as from regions 1.5 and 3 cm above the lesions. Within necrotic lesions, bacteria were present at high levels and were primarily localized in the intracellular regions of the tomato pith, particularly surrounding dead or necrotic cells (Fig. 3). At 1.5 cm above the necrotic lesion, both bacteria were detected at substantially lower levels and were localized in both intracellular pith regions and vascular bundles (Fig. 3). Bacteria were not detected 3 cm above the lesion in most samples; in the few positive samples, they were mainly confined to single vascular bundles (Fig. 3). Notably, we initially labeled Pmed G51 with GFP as well. However, while GFP-fluorescent kanamycin-resistant transformants were initially recovered following transformation, most clones gradually lost detectable GFP fluorescence over time. Consequently, GFP fluorescence was not detectable in tomato tissues at 14 dpi, despite high recovery of kanamycin-resistant bacteria from the inoculation site. Due to this instability of GFP expression, Pmed localization assays were excluded from the study.

**Fig. 3.**
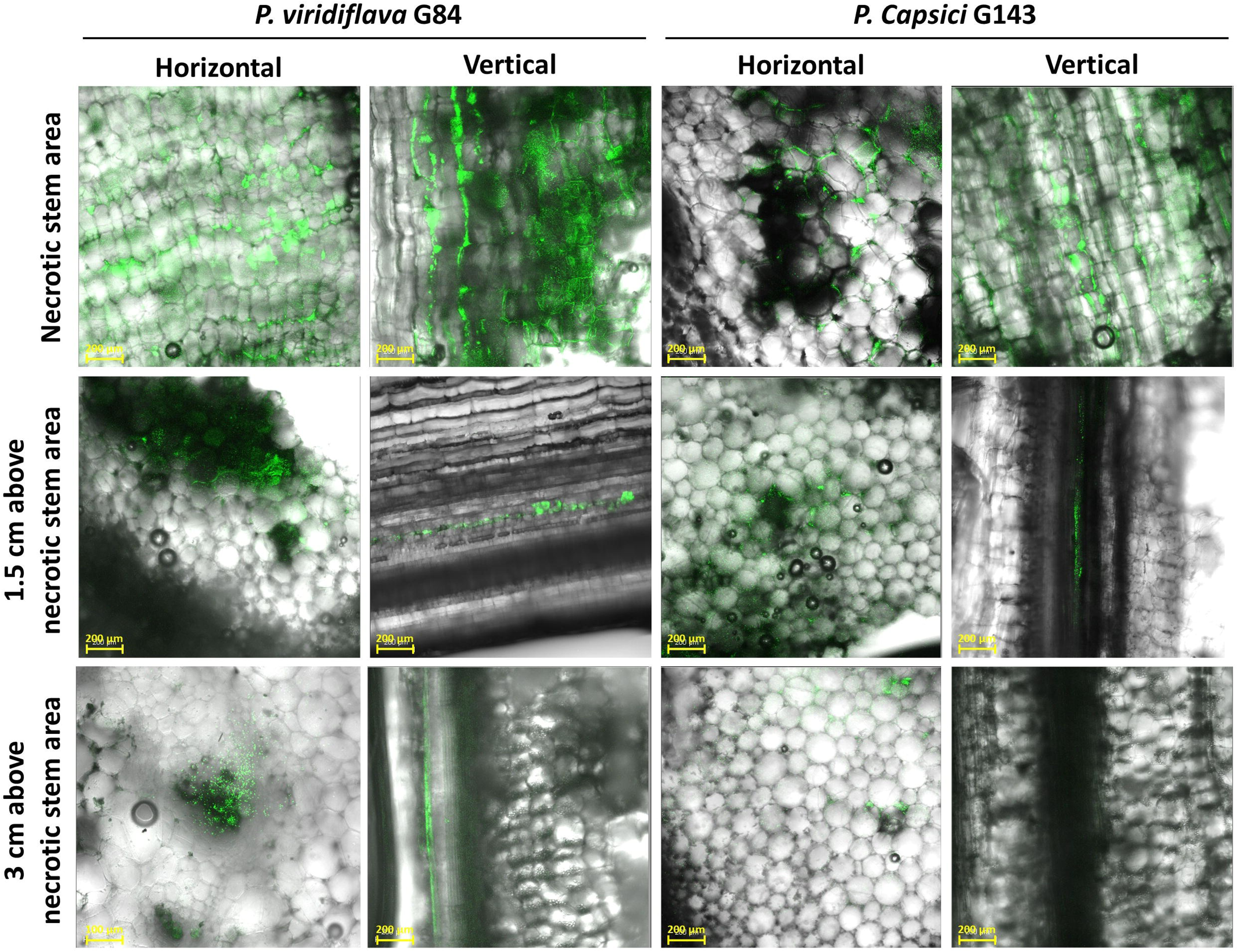
In situ localization of pith necrosis–associated bacteria during infection. Tomato cv. Ikram plants were inoculated with GFP-labeled *P. viridiflava* G84 or *P. capsici* G143. Bacterial colonization within the stem was visualized by confocal laser scanning microscopy using a GFP filter at 14 days post inoculation, at the margins of necrotic stem tissue and in non-symptomatic tissue located 1.5 and 3 cm above the necrotic areas.

These results indicate that only a subset of pith necrosis–associated bacterial species possesses consistent pathogenic potential in tomato and that disease severity is associated with limited but spatially structured bacterial colonization, characterized by high bacterial densities within necrotic pith tissue and restricted upward movement, primarily through vascular bundles.

### Viral infection exacerbates tomato pith necrosis systems

Pith necrosis is traditionally classified as a disease caused by weak opportunistic pathogens, which is consistent with the high plant-to-plant variability observed in symptom severity. Given that pith necrosis has become extremely prevalent in Israel over the past decade, coinciding with the introduction of ToBRFV and PepMV, we hypothesized that viral infection may contribute to or exacerbate disease development. To test this hypothesis, we conducted co-inoculation experiments using pith necrosis–associated *Pseudomonas* isolates (Pmed, Pvir, and Pcap) together with ToBRFV, PepMV, or both viruses, and monitored lesion size and bacterial populations. Given that lesion expansion patterns were highly variable among plants (Fig. S2), we applied an unbiased approach to quantify lesion area using image analysis with ImageJ. Infection with ToBRFV and PepMV, with or without *Pseudomonas* inoculation, significantly affected plant growth and development (Fig. S3). Co-inoculation of each *Pseudomonas* isolate with both ToBRFV and PepMV resulted in a significant 40–100% increase in lesion size compared with bacterial inoculation alone (Fig. 4A, B). In contrast, co-inoculation with each virus individually produced a similar but less consistent trend, with a statistically significant increase observed only when either virus was co-inoculated with Pvir (Fig. S4). We next assessed whether viral infection affected local and distal bacterial colonization by bacterial isolation and CFU quantification. Consistent with the GFP localization data, distal bacterial populations measured 2 cm above the inoculation site were reduced by ∼500-fold relative to populations at the infection site (Fig. 4C). Viral infection resulted in either no effect or only a minor (10–20%) increase in bacterial colonization, which was statistically significant only for distal colonization by Pmed (Fig. 4C). Overall, these results suggest that viral infection can intensify disease symptoms without substantially increasing bacterial spread, highlighting a potential role of ToBRFV and PepMV in pith necrosis severity.

**Fig. 4.**
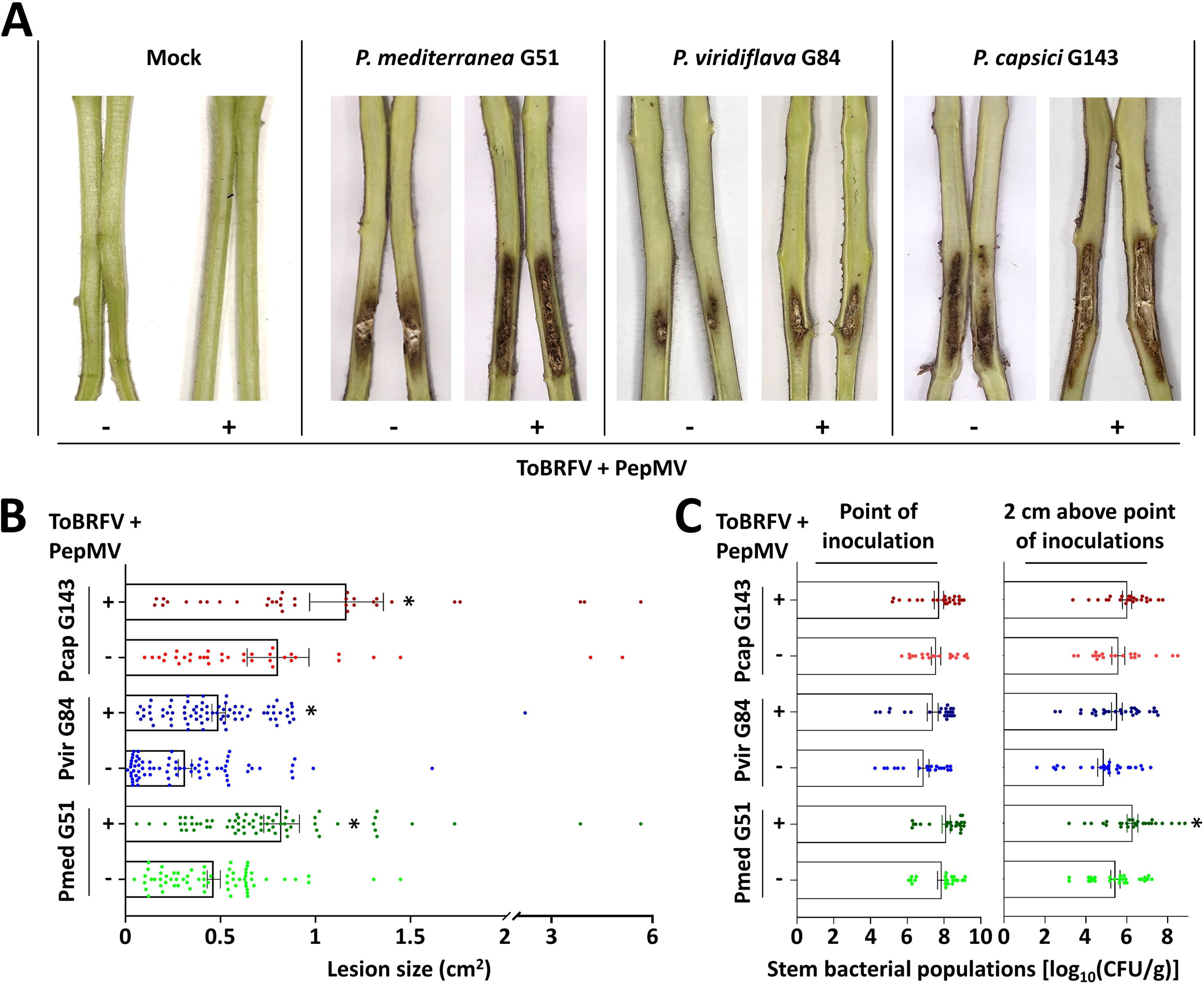
Effect of combined ToBRFV and PepMV infection on lesion size induced by pith necrosis–associated bacteria. Tomato cv. Ikram plants were co-inoculated with the pith necrosis–associated bacteria *P. mediterranea* G51 (Pmed), *P. viridiflava* G84 (Pvir), *P. capsici* G143 (Pcap), or a no-bacteria control (mock), together with ToBRFV and PepMV (+) or a no-virus control (−). **(A)** Representative split stems were photographed 30 days post inoculation (dpi). **(B)** Pith necrosis lesion size was quantified using ImageJ at 30 dpi. Bar graphs depict the mean values, standard errors, and individual data points from at least 35 biological replicates pooled from five independent experiments, each composed of six to 17 plants. **(C)** Bacterial populations were quantified at 30 dpi at the point of inoculation and 2 cm above the inoculation site. Bar graphs depict the mean values, standard errors, and individual data points from at least 20 biological replicates pooled from four independent experiments each containing 3-6 plants. **(B, C)** Asterisks indicate statistically significant differences (Mann–Whitney U test, *p* < 0.05) between ToBRFV and PepMV infected plants and the corresponding no-virus controls inoculated with the same bacterial strain.

### Viral infection attenuates tomato bacterial canker symptoms

Given that viral infection enhanced pith necrosis symptoms, we tested whether a similar phenomenon could be observed for bacterial canker, for which co-infections were also detected in greenhouses in the northwestern Negev. ‘Ikram’ tomato plants were co-inoculated with *C. michiganensis* together with ToBRFV and/or PepMV and monitored for wilt symptoms as well as local and distal bacterial populations. In contrast to our observations for pith necrosis, tomato plants inoculated with ToBRFV and/or PepMV exhibited a reduction in wilt symptoms, which were apparent in all *C*. *michiganensis* -inoculated plants but were significantly less severe in virus-infected plants compared with the virus-free control (Fig. 5A, 5B, and S5). Assessment of local and distal stem bacterial populations revealed no differences in colonization capacity among treatments, with similar bacterial levels observed in virus-infected and virus-free plants (Fig. 5C). These findings indicate that viral infection can modulate disease symptom development in bacterial canker without affecting bacterial colonization.

**Fig. 5.**
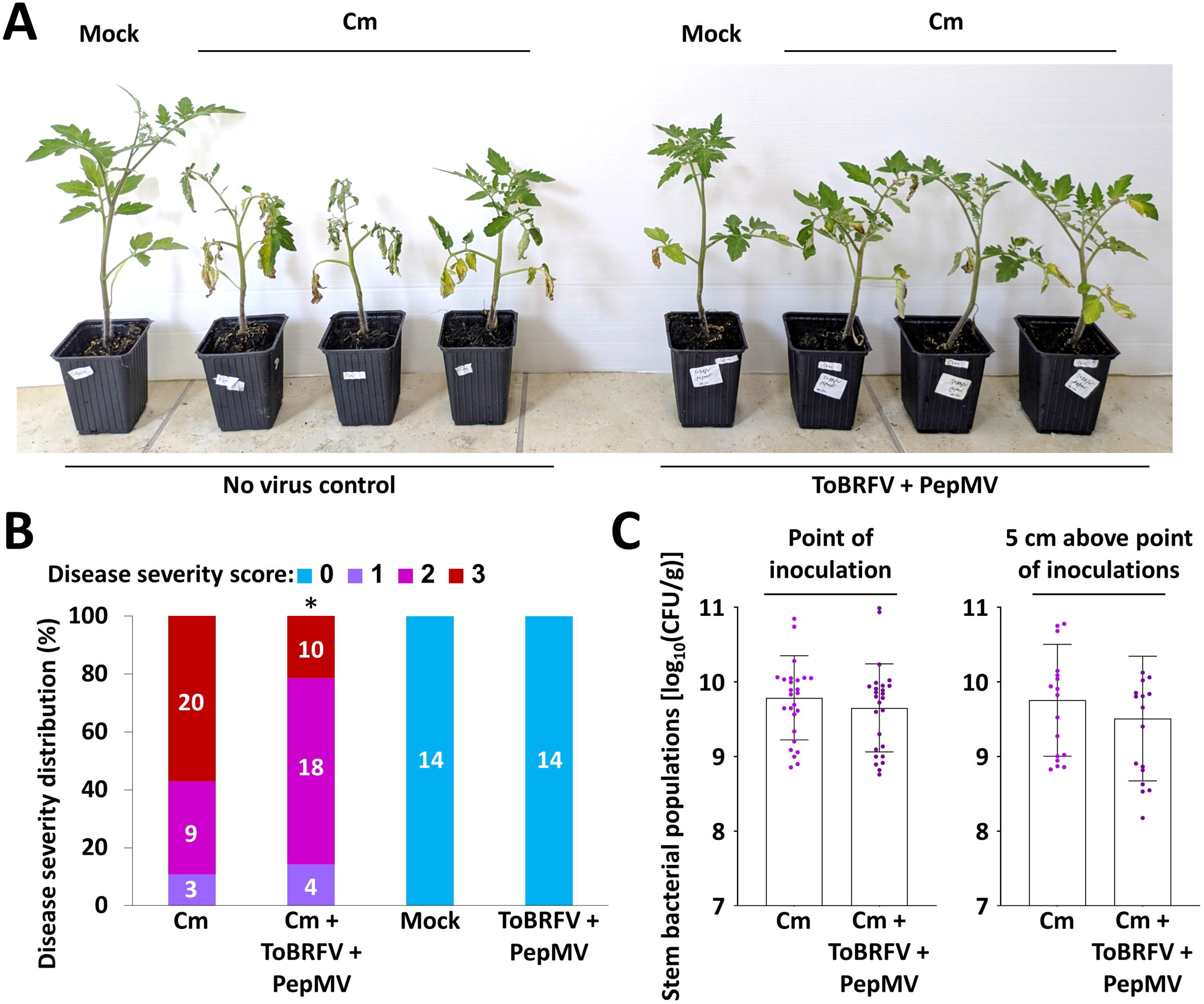
Effect of combined ToBRFV and PepMV infection on wilt symptoms induced by *Clavibacter michiganensis*. Tomato cv. Ikram plants were co-inoculated with *Clavibacter michiganensis* (Cm) NCPPB382 or a no-bacteria control (mock), together with ToBRFV and PepMV or a no-virus control. (**A**) Representative plants were photographed at 14 days post inoculations (dpi). (**B**) Wilt symptoms were scored at 14 dpi by the percentage of wilted leaves according to the following scale: 0 – no wilting, 1 = 1-25%, 2= 26-60%, 3= 61-100%. The graph depicts the distribution of 32 repeats for each clone pooled from five experiments each containing 6-10 plants. Number of plants with each score are embedded within the bar. Asterisks indicate significant difference (chi-squared test, *p* < 0.05) between ToBRFV and PepMV infected plants and the corresponding no-virus control. (**C**) Bacterial populations were quantified at 14 dpi at the point of inoculation and 5 cm above the inoculation site. Bar graphs depict the mean values, standard errors, and individual data points from at least 18 biological replicates pooled from three independent experiments each containing 6-10 plants.

## Discussions

Tomato is one of the most economically significant vegetable crops grown worldwide and according to the FAO 2023 the tomato production in 2023 reached a record high, with estimates exceeding 192 million metric tons [28]. In plant science research, tomato serves as a premier model system for studying plant-pathogen interactions due to the extensive genetic tools available [29]. However, tomato production is severely constrained by a diverse array of viral, bacterial, and fungal diseases, with outbreaks of ToBRFV, bacterial wilt, and bacterial canker posing substantial threats to regional industries [1]. While diseases are typically studied in isolation under controlled conditions, such approaches often overlook the complex pathogen-to-pathogen interactions that influence disease etiology and crop yield. This study employs a more holistic approach by surveying field-scale co-infections and assessing the synergistic effects of these interactions on disease progression. Our findings provide new insights into how primary viral infections significantly alter the severity of bacterial diseases. We show that pith necrosis and bacterial canker outbreaks in Israeli tomato production occur in the presence of ToBRFV and PepMV infections, which have become endemic in the country. We demonstrated that viral infection differentially alters disease severity, resulting in the exacerbation of pith necrosis symptoms and the attenuation of bacterial canker symptoms. In addition, this study identified the causal and associated agents of pith necrosis in Israel and demonstrated that the disease can be caused by different pathogens. Notably, in numerous cases, bacteria that are heavily associated with the disease in the field cannot cause it in controlled laboratory experiments.

Tomato pith necrosis has been primarily associated with *P. corrugata*, *P. mediterranea*, and members of the *P. fluorescens* group and the *P. syringae* species complex, as well as SRPs, for which the disease is sometimes classified separately as stem soft rot [16,18,19,30,31]. In addition, Disease outbreaks in Italy have attributed tomato pith necrosis to *X. e*. pv. *perforans*, possibly involving a bacterial complex with soil-borne *Pseudomonas* spp. [17]. Notably, many published studies regarding tomato pith necrosis are descriptive case reports that focus primarily on bacterial isolation and identification, without providing experimental evidence, such as pathogenicity assays, to conclusively demonstrate a causal role for the isolated bacteria in disease development. Following isolation from grower-supplied tomato plants exhibiting pith necrosis-like symptoms in Israel, we found that the pith necrosis phenotype was associated with a diverse bacterial community, many members of which have previously been implicated in the disease in other regions in the world. However, a substantial proportion of the isolates failed to induce disease symptoms in controlled laboratory inoculation assays. These results suggest that such isolates are may not be the primary causal agents, but may instead act as opportunistic colonizers following tissue maceration triggered by biotic or abiotic factors that may exacerbate the phenomenon, or cause disease only under specific greenhouse conditions that were not reproduced in our experimental system. Our analysis identified only four pith necrosis–associated bacterial species that were able to cause pith necrosis disease following artificial inoculation. These include *P. mediterranea*, the traditionally recognized causal agent of pith necrosis, *P. capsici*, *X. e.* pv*. perforans*, and *P. viridiflava*. For all four pathogens, infection assays resulted in highly variable disease severity among plants maintained under similar conditions. We note that the inoculation method used for pith necrosis, which relied on direct transfer from bacterial plates, may introduce higher variability due to the lack of standardized inoculum concentration. However, we selected this approach because inoculation using liquid suspensions of resuspended bacteria standardized to the same OD (as used for *C. michiganensis*) was inconsistent and in some cases failed to produce reliable infections, resulting in even greater variability in symptom severity.

Given that the increased incidence of vascular collapse in Israeli greenhouse tomatoes coincided with the emergence of ToBRFV and the widespread increase in PepMV infections, which have overtaken the national tomato industry [7], we hypothesized that this increase results from enhanced host susceptibility caused by viral infection. Our data partially support this hypothesis, showing that viral infection enhanced disease caused by pith necrosis–associated bacteria but attenuated bacterial canker disease. Despite the clear increase in lesion size in the background of viral infection, the symptoms observed under experimental conditions were significantly milder than those reported in commercial greenhouse plots. Bacteria-induced necrosis following artificial inoculation was largely internal and did not manifest as vascular collapse or external lesions. This discrepancy may be due to glasshouse conditions that do not fully recapitulate commercial greenhouse environments, including differences in irrigation regimes, humidity, temperature, and plant density, and/or the involvement of additional contributing factors [32]. Notably, pith necrosis has been reported to be associated with improper fertilization and excessive nitrogen availability [15]. It is possible that, in response to yield losses caused by ToBRFV and PepMV, growers adjusted agricultural practices, such as fertilization, pruning, or pesticide use, which may inadvertently influence plant physiology and increase susceptibility to pith necrosis. However, this remains speculative and was not directly tested in this study. Future studies should assess pith necrosis under controlled variations in nitrogen and irrigation in both virus-infected and non-infected plants. Replicated trials that better reflect commercial greenhouse conditions would further help disentangle the contributions of viral infection and agronomic practices to disease severity.

[33–35]ToBRFV and PepMV infections in tomato differentially modulated bacterial disease severity, increasing pith necrosis while reducing bacterial canker. However, the molecular and physiological mechanisms underlying this contrasting effect remain unclear. Viral infections induce extensive transcriptional reprogramming, leading to profound developmental and physiological changes at the cellular, tissue, and whole-plant levels, as well as shifts in secondary metabolism and phytohormone homeostasis [36]. These diverse effects have been reported to influence host responses to both biotic and abiotic stresses. Multiple studies have linked viral infection to increased tolerance to heat or drought in various plant systems [37–39], presumably due to stress priming mediated by activation of host immune responses [40]. In contrast, the effect of viral infection on susceptibility to microbial pathogens varies widely, depending on the host, virus, and pathogen. For example, rice necrosis mosaic virus (*Bymovirus oryzae*) increases rice susceptibility to bacterial, fungal, and viral pathogens [43], and cucumber green mottle mosaic virus (*Tobamovirus viridimaculae*) increases susceptibility to *Pythium* in cucumber [41]. Conversely, potato virus X (*Potexvirus ecspotati*) and plum pox virus (*Potyvirus plumpoxi*) can enhance tolerance to *Pseudomonas syringae* [42], while tomato mosaic virus (*Tobamovirus tomatotessellati*) reduces susceptibility to bacterial spot and speck in tomato [43]. Similarly, ToBRFV has been shown to promote PepMV spread [7], and can differentially affect pathogen interactions, increasing susceptibility to *Botrytis cinerea* while reducing susceptibility to *Xanthomonas euvesicatoria* [44].

It has been suggested that salicylic acid (SA) accumulation during viral infection, reported for both ToBRFV and PepMV [44,45], may influence host responses to microbial pathogens, potentially promoting susceptibility to necrotrophs while reducing susceptibility to hemibiotrophs. Although tomato pith necrosis has not been extensively studied in the context of phytohormone signaling, its opportunistic nature and the strong association between bacterial presence and tissue necrosis suggest a more necrotrophic-like behavior. In this context, viral-induced SA accumulation could plausibly exacerbate tissue necrosis, leading to larger lesions. In contrast, ToBRFV and PepMV attenuated bacterial canker symptoms caused by *C. michiganensis*. As a highly specialized vascular pathogen with a hemibiotrophic lifestyle, *C. michiganensis* may be affected differently by host immune priming associated with viral infection [11]. Viral infection could therefore either enhance resistance to *C. michiganensis*, as previously reported for SA-mediated responses [46], or reduce wilt symptoms that overlap with drought-related physiological stress. Notably, although still endemic throughout the country, *C. michiganensis* is currently considered a much less significant pest compared to the severe outbreaks reported from the 2000s to the mid-2010s. Further studies are warranted to determine whether the widespread occurrence of ToBRFV and PepMV is associated with this change in disease prevalence.

This study highlights the importance of considering multi-pathogen interactions when evaluating disease development in crop systems, as primary viral infections can fundamentally reshape host–microbe relationships and disease outcomes. Our findings suggest that the increasing prevalence of endemic viral pathogens may have unanticipated consequences for the epidemiology and severity of bacterial diseases in tomato, contributing both to disease exacerbation and attenuation depending on pathogen lifestyle. Incorporating such interactions into disease management and experimental frameworks will be essential for accurately predicting disease risk and improving the resilience of intensive tomato production systems.

## Acknowledgment

We would like to thank Dr. Shulamit Manulis-Sasson (Volcani Institute) for providing access to her collection of pith necrosis–associated bacteria and associated metadata from 2014–2015. This research was supported by Chief Scientist of the Israeli Ministry of Agriculture, grant numbers 20-02-0166 and 20-02-0208

## Supporting information

**Table S1.** Primers used during this study

**Fig. S1.** Symptoms caused by artificial inoculation of tomato with *Pectobacterium aroidearum*.

**Fig. S2.** Variation in symptom severity caused by pith necrosis–associated bacteria.

**Fig. S3.** Effect of ToBRFV and PepMV on plant development.

**Fig. S4.** Effect of ToBRFV or PepMV infection on lesion size induced by pith necrosis–associated bacteria.

**Fig. S5.** Effect of ToBRFV or PepMV infection on n wilt symptoms induced by *Clavibacter michiganensis*.

