## Supplementary Figures and tables for "Tomato brown rugose fruit virus and pepino mosaic virus differentially modulate disease severity of bacterial pith necrosis and bacterial canker in tomato"

**Table S1. Primers used during this study**

| **Name** | **Gene target** | **Bacteria** | **F primer** | **R primer** |
| --- | --- | --- | --- | --- |
| Uni16S | 16S rDNA | All bacteria | AGAGTTTGATCCTGGCTCAG | GGTTACCTTGTTACGACTT |
| rpoBPs | *rpoB* | Pseudomonadales order | GACAAGATGGCCGGTCGTCACGGTAAC | TTCGGTTTCCAGATCGATATCGAT |
| recAPs | *recA* | Pseudomonadales order | GCCTTGGCTGCGGCCTTGGGTCAGATCG | TAGGCATACCAGGCACCGGACTTCTCGA |
| gyrBPs | *gyrB* | Pseudomonadales order | CGGAAGAAGAAGGTCAGCAGCAGGGTACGGAT | GCACCGCGATCATGCCTTCGC |
| gyrBXan | *gyrB* | Xanthomonadales order | GAGCCGCAGTACCCGCTCAAGC | GCACCGCGATCATGCCTTCGC |
| gyrBEn | *gyrB* | Enterobacterales order | CCATCGACGTCAGCATCGGTCATGATGAT | TGCAGTGGAACGATGGTTTCCAGGAAAA |
| gyrBMic | *gyrB* | Micrococcales order | TCCAGCAGATGGCCTTCCTCAAC | TGCGGCTCGCCGAGCTTCACGG |
| chpCdi | *chpC* | *Clavibacter michiganensis* | TTGGAGTGCAGTGTTCTTCG | TACACCATCGTGCTCTGCTC |

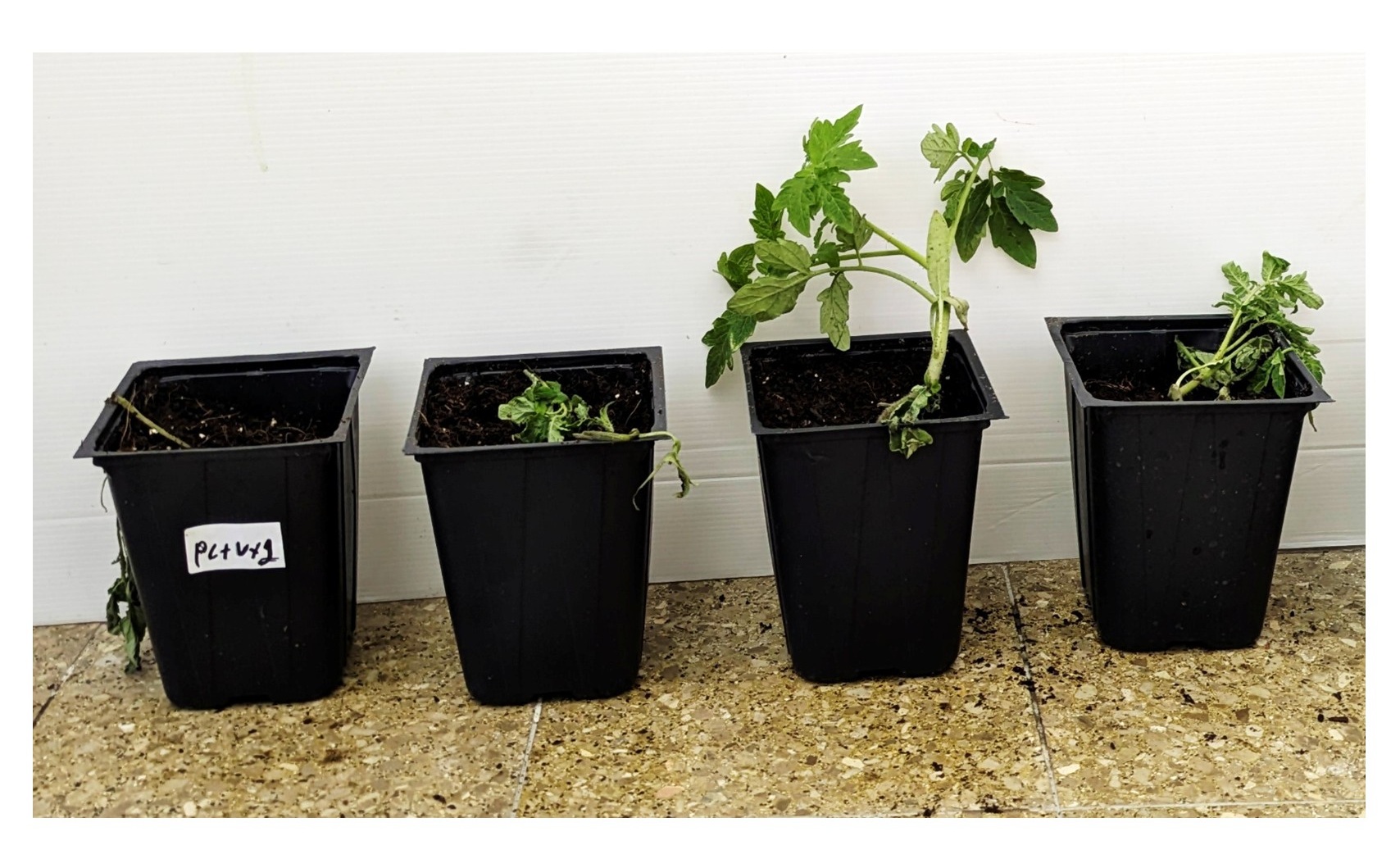

**Fig. S1. Symptoms caused by artificial inoculation of tomato with Pectobacterium aroidearum.** Tomato cv. Ikram plants were inoculated with the pith isolate P. aroidearum G201 by puncturing the stem between the cotyledons using a toothpick soaked in a bacterial suspension (OD600 = 0.1). Images were taken 72 h post-inoculation.

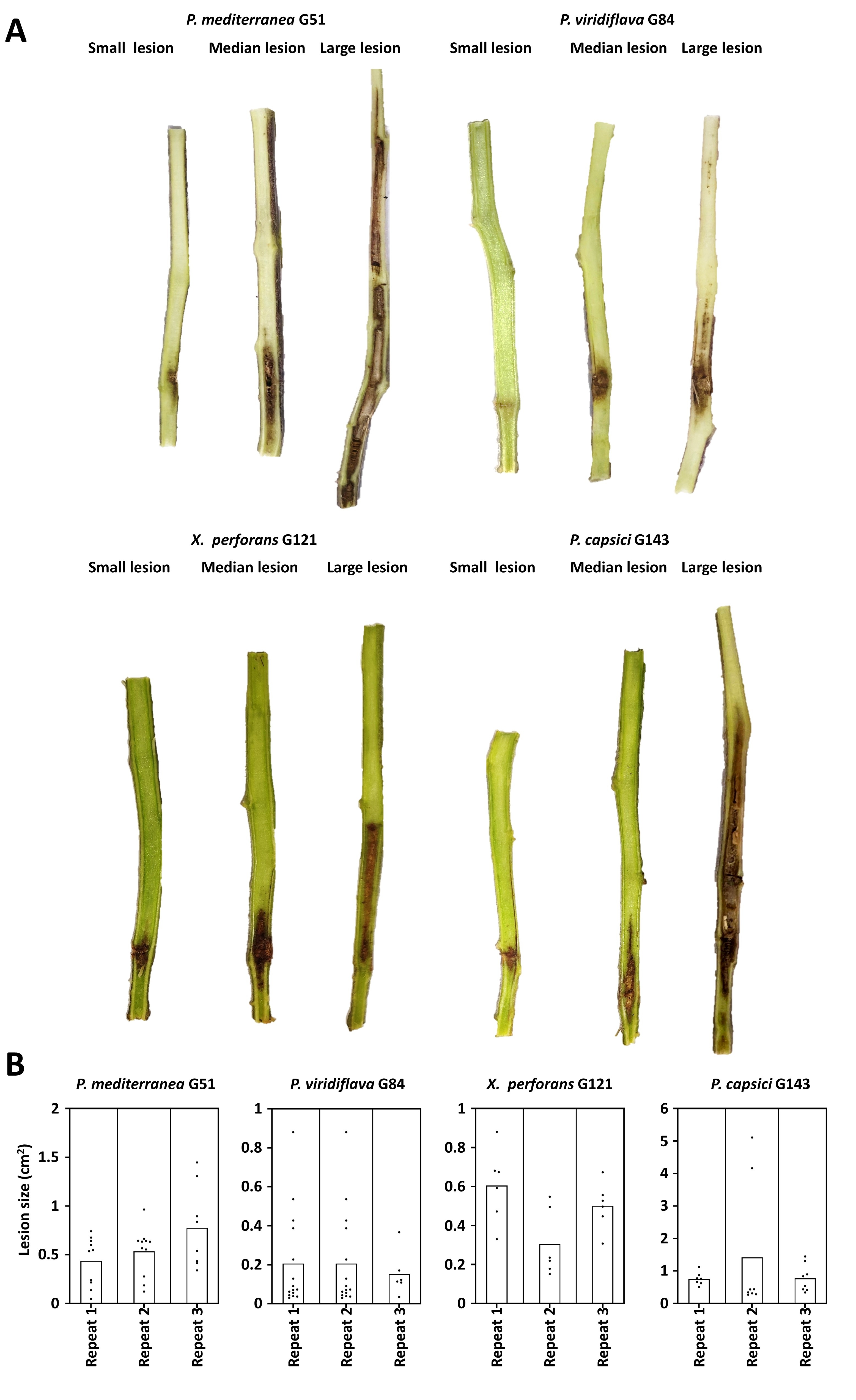

**Fig. S2. Variation in symptom severity caused by pith necrosis–associated bacteria.** Tomato cv. Ikram plants were inoculated with the pith necrosis–associated bacteria P. mediterranea G51, P. viridiflava G84, X. e. perforans G121, or P. capsici G143. (**A**) Representative images showing different levels of symptom severity were taken 30 days post inoculation. (**B**) Pith necrosis lesion size was quantified at 30 dpi using ImageJ. Bar graphs show mean values, with individual data points representing biological replicates from three independent experiments per pathogen.

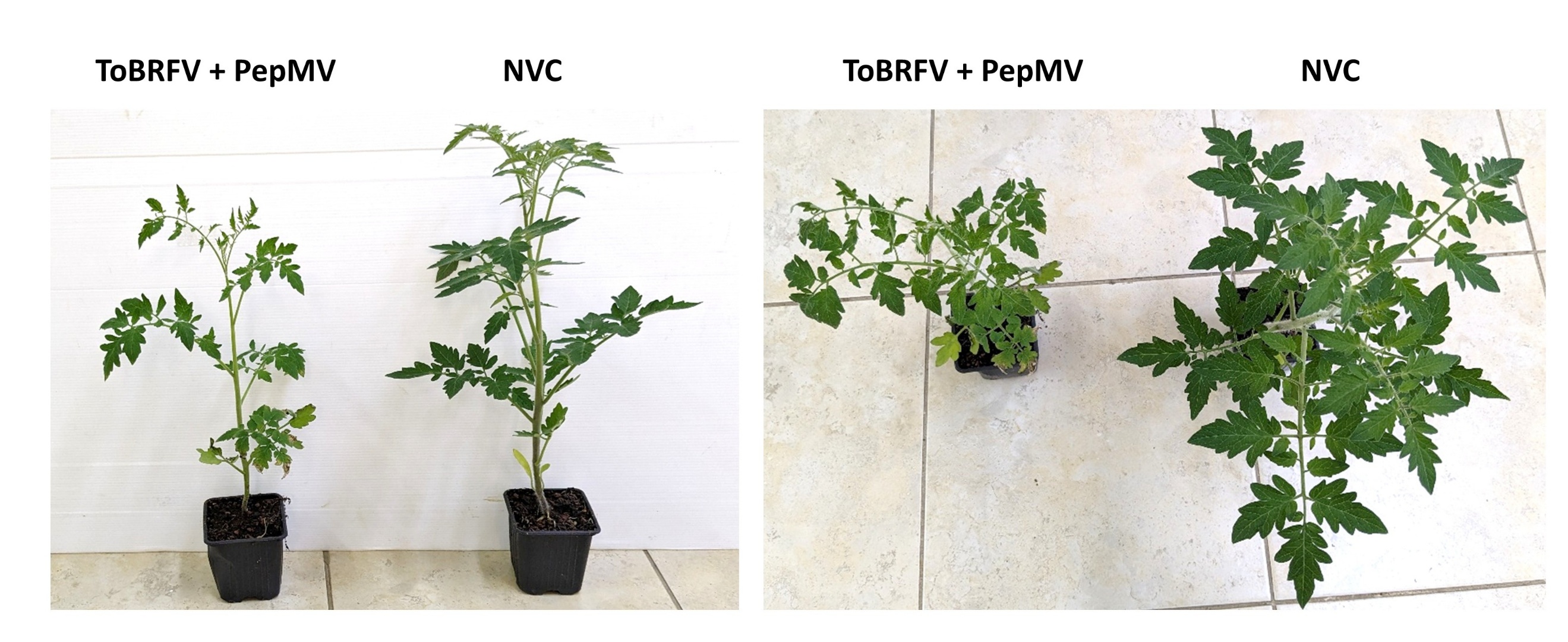

**Fig. S3. Effect of ToBRFV and PepMV on plant development**. Four-leaf–stage tomato cv. Ikram plants were inoculated with a combination of ToBRFV and PepMV. Plants were photographed 30 days post inoculation.

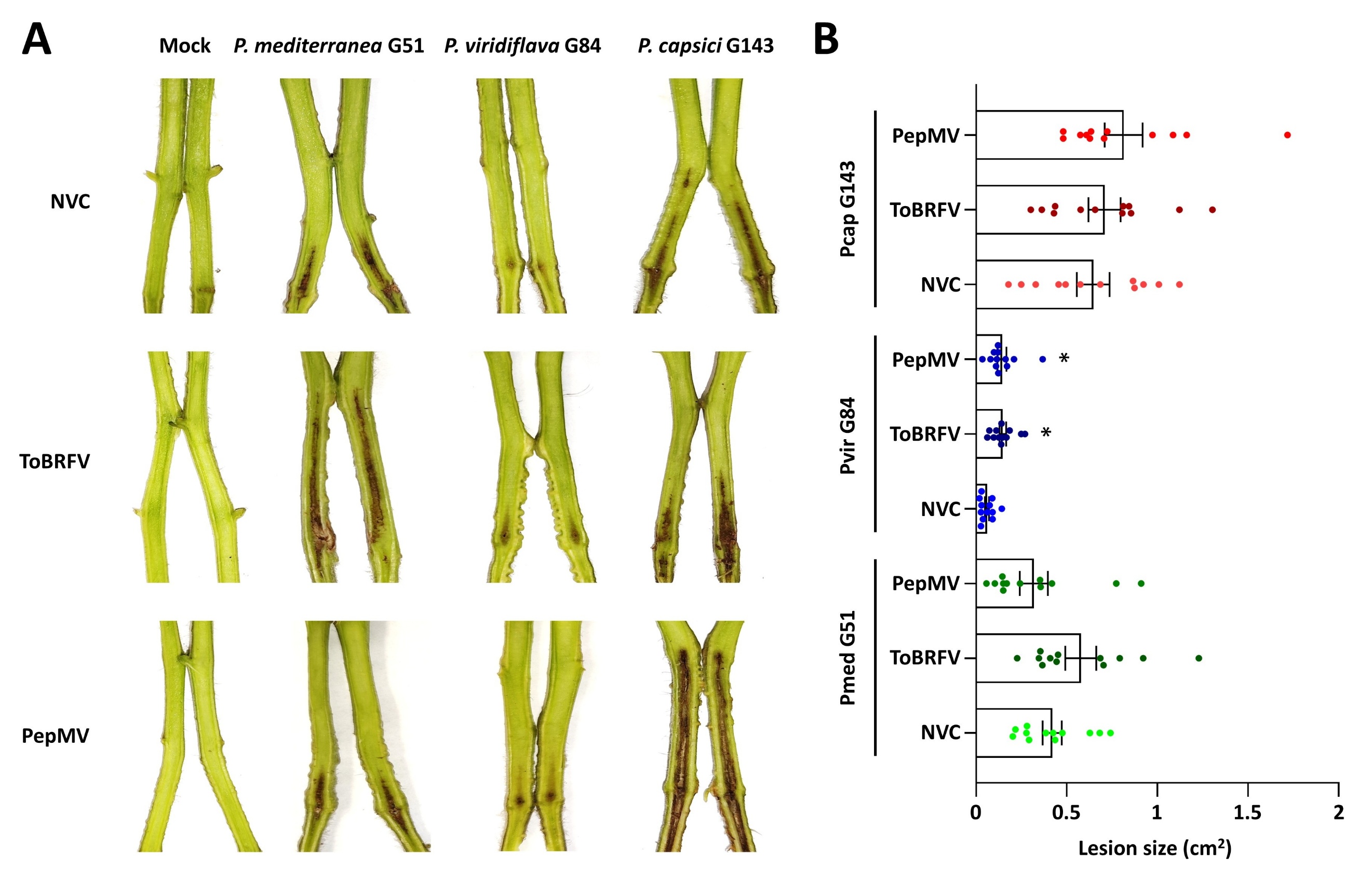

**Fig. S4. Effect of ToBRFV or PepMV infection on lesion size induced by pith necrosis–associated bacteria.** Tomato cv. Ikram plants were co-inoculated with the pith necrosis–associated bacteria P. mediterranea G51 (Pmed), P. viridiflava G84 (Pvir), P. capsici G143 (Pcap), or a no-bacteria control (mock), together with ToBRFV, PepMV (+) or a no-virus control (NVC). **(A)** Representative split stems were photographed 30 days post inoculation (dpi). **(B)** Pith necrosis lesion size was quantified using ImageJ at 30 dpi. Bar graphs depict the mean values, standard errors, and individual data points from 12 biological replicates pooled from two independent experiments. Asterisks indicate statistically significant differences (Mann–Whitney U test, p < 0.05) between ToBRFV or PepMV infected plants and the corresponding NVCs inoculated with the same bacterial strain.

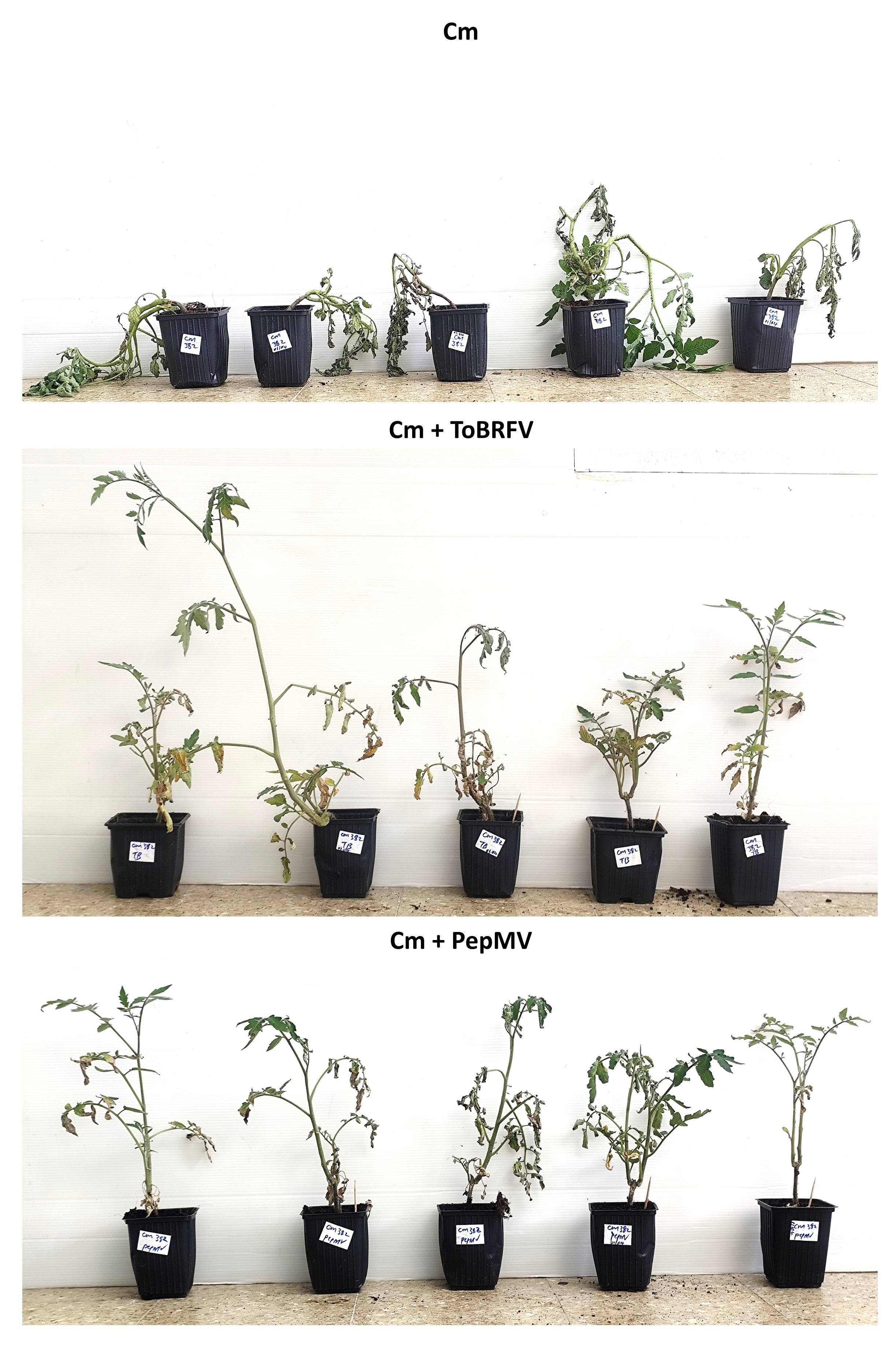

**Fig. S5. Effect of ToBRFV or PepMV infection on n wilt symptoms induced by *Clavibacter michiganensis*.** Tomato cv. Ikram plants were co-inoculated with *Clavibacter michiganensis* (Cm) NCPPB382 or a no-bacteria control (mock), together with ToBRFV, PepMV, or a no-virus control. Representative pictures were taken at 14 days post inoculations.
